# Aggregation-mediated microcolony shields bacteria from contact-dependent competition and maintains phenotypic heterogeneity

**DOI:** 10.64898/2026.03.26.714566

**Authors:** Margot Marie Dessartine, Julien F. Giraud, Artemis Kosta, Guoliang Qian, Éric Cascales, Jean-Philippe Côté

**Affiliations:** Département de biologie, Faculté des sciences, Université de Sherbrooke, Sherbrooke, QC J1K 2R1, Canada; Laboratoire d’Ingénierie des Systèmes Macromoléculaires (LISM), Aix-Marseille Université, CNRS, UMR7255, Marseille, France; Service de microscopie, Institut de Microbiologie de la Méditerranée, Aix-Marseille Université, CNRS, UAR2044, Marseille, France; State Key Laboratory of Agricultural and Forestry Biosecurity, College of Plant Protection, Nanjing Agricultural University, Nanjing 210095, P.R. China

**Keywords:** bacterial competition, secretion system, microcolony formation, adhesive structure, phenotypic diversity, antagonism, fimbriae, type VI secretion, type IV secretion, contact-dependent growth inhibition, colicins, bacteriocins, antibiotics

## Abstract

Bacteria are adaptable microorganisms capable of surviving in a wide variety of environments. One common strategy for persistence and attachment to surfaces is the formation of microcolonies, often controlled by cell surfaces structures that mediate adhesion such as fimbriae. In this study, we show that type 1 fimbriae, and other adhesins structures such as autotransporter adhesins, trigger microcolony formation which confer resistance against a broad range of short-range contact-dependent weapons (e.g. T6SS, T4SS, CDI) but not against long-range diffusible weapons (e.g. diffusible toxins like colicins). Interestingly, we identify “sheltered” bacteria within microcolonies that benefit from collective protection without expressing the adhesive structure. Our findings demonstrate that adhesive structures not only improve survival in hostile environments by promoting microcolony formation but also maintain phenotypic heterogeneity within the microcolony, highlighting the importance of social behavior in bacterial adaptation to changing environments.

## Introduction

From the synchronized swimming of fish schools to the neural networks enabling cognition, collective behaviors are a fundamental feature in living organisms. In bacteria, two key parameters influencing collective behaviors are cell density and spatial localization ^1,2^. Bacterial coordination and interaction can be facilitated by aggregation, a process that promotes the formation of microcolonies. Cell clustering is mediated by cell surface adhesins and extracellular adhesive appendages such as fimbriae and pili ^3–5^, and plays an important role in the formation of biofilms, the attachment to biotic and abiotic surfaces, and the colonization of ecological niches ^6–9^. Microcolonies can also provide protection against external threats such as host immune responses and environmental stresses (e.g., antibiotic exposure) ^8,10,11^.

In a previous study, we developed a high-throughput interbacterial competition assay to identify *Escherichia coli* K12 mutants from the Keio collection ^12^ that were resistant to contact-dependent type VI secretion system (T6SS) attacks ^13^. This screen identified a *fimE* mutant that exhibited resistance to *Cronobacter malonaticus* T6SS assaults. In a Δ*fimE* mutant, type I fimbriae were overexpressed compared to the parental strain, promoting cell aggregation, microcolony formation and collective protection against T6SS ^13^. These observations raised broader questions about the role of microcolonies in bacterial resistance and microbial interaction, notably whether microcolony formation may represent a nonspecific defense strategy and whether microcolonies can serve as shelters for non-aggregating bacteria.

Here, we show that type 1 fimbriae-mediated microcolonies resist assaults from various T6SS^+^ species such as *C. malonaticus*, enteroaggregative *Escherichia coli* (EAEC) and *Citrobacter rodentium*. Interestingly, the protection provided by the microcolonies is not restricted to T6SS, as they also protected against other contact-dependent antibacterial competition mechanisms, such as the type IV secretion (T4SS) and contact-dependent growth inhibition (CDI) systems. However, microcolonies are susceptible to diffusible chemicals and toxins such as hydrogen peroxide and colicins. Finally, we show that bacteria unable to produce type 1 fimbriae can be sheltered within microcolonies, where they benefit from assaults by competitors, hence promoting phenotypic heterogeneity within bacterial population. We further demonstrate that other factors promoting aggregation, such as autotransporter adhesins secreted by the type V secretion systems (T5SS), promote microcolony formation and confer protection against T6SS assaults. These findings highlight the protective and social functions of microcolonies, providing insight into bacterial survival strategies under competitive and hostile conditions.

## Results

### Engineering of *E. coli* locked *fimS* promoter strains

We previously showed that type 1 fimbriae-dependent microcolonies conferred resistance against T6SS attacks from *C. malonaticus* strain 3267 (CM3267) ^13^. Expression of the *fim* operon that encodes type 1 fimbriae is controlled by a phase variation mechanism involving inversion of the *fimS* promoter by the FimB and FimE recombinases ^14,15^. In our previous work, we observed that the deletion of the *fimE* gene, the recombinase that preferentially switches the promoter from the ON- to the OFF-phase, led to an increase in fimbriae production and a partial resistance to T6SS. However, a significant number of Δ*fimE* cells remain in the OFF phase. Indeed, only 50% of Δ*fimE* cells exhibited type 1 fimbriae at their bacterial cell surface. To study the process of microcolony-mediated resistance with homogeneous populations, we engineered two *E. coli* strains with the *fimS* promoter locked in either the ON (locked-ON) or OFF (locked-OFF) orientation, thereby preventing phase variation. For this, one of the 9-bp inverted repeats flanking the 314-pb invertible *fimS* promoter was deleted to prevent the permutation.

Negative-stain electron microscopy confirmed that 100% of locked-ON cells produced type 1 fimbriae, whereas none of the locked-OFF cells did (Figure 1A and 1C). Production of type 1 fimbriae correlated with the formation of microcolonies, as shown by confocal microscopy under bright-light field (Figure 1B). Type 1 fimbriae are tipped by the FimH adhesin, which binds mannosylated receptors on various eukaryotic cells, promoting their agglutination. To confirm that the aggregation properties were mediated by type 1 fimbriae, wild-type (WT)-like *E. coli* BW25113 Δ*yejO* cells ^12,13^, locked-ON and locked-OFF cells were mixed with *Saccharomyces cerevisiae* cells and observed. As previously reported, WT-like and locked-OFF cells did not induce yeast agglutination while yeast cells incubated with locked-ON cells formed visible aggregates (Figure 1D). Addition of the competitive inhibitor D-mannose inhibited FimH-mediated agglutination of yeast cells in all cases (Figure 1D). Growth analyses revealed a slight decrease in growth rate of the locked-ON strain, likely due to the fimbriae overexpression (Figure 1E). Taken together, these results confirmed the homogeneity of the two engineered strains in terms of fimbriae production and cell aggregation.

**Figure 1.**
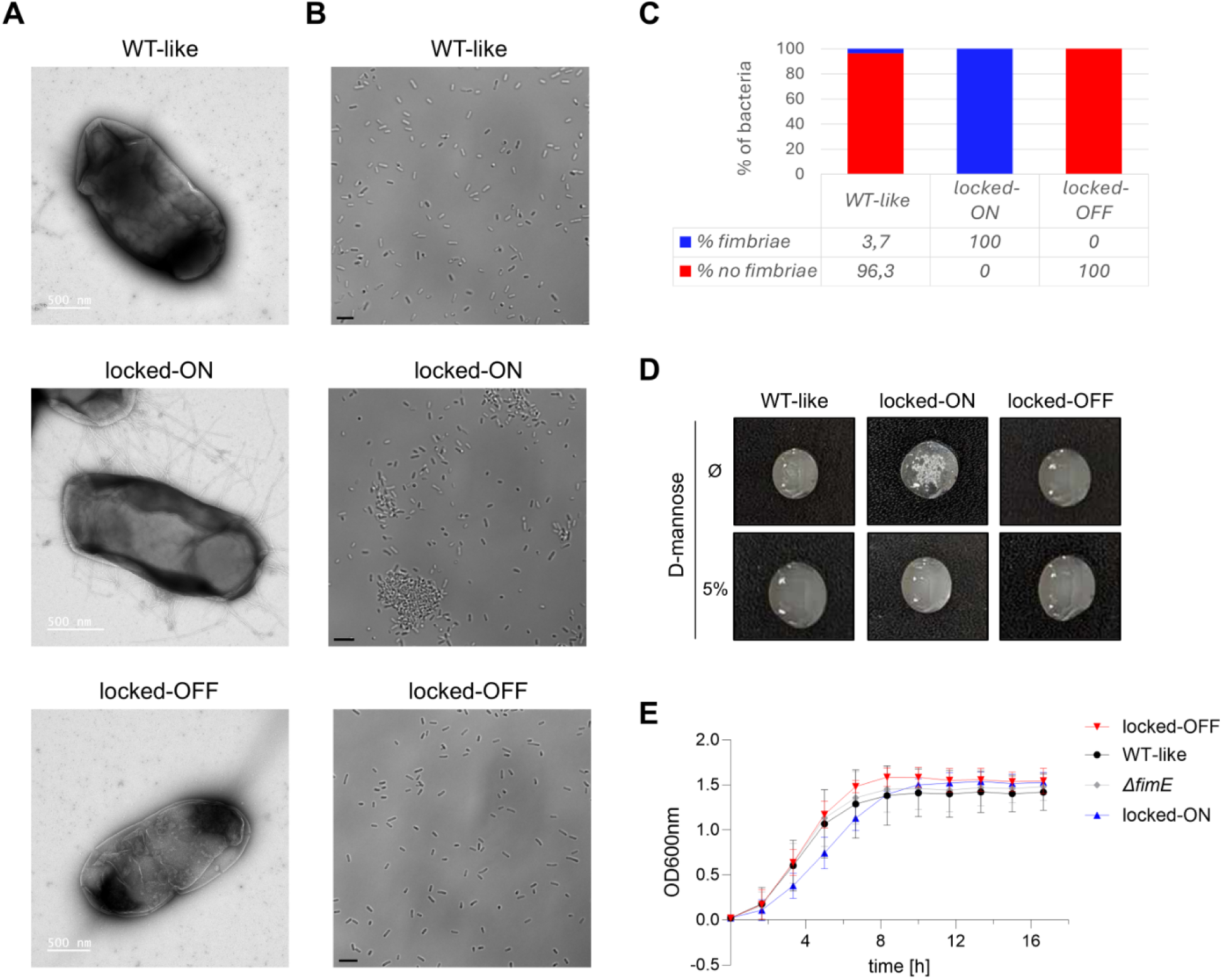
Phenotypic analysis of *E. coli* locked-ON and locked-OFF *fimS* promoter mutants. Representative negative-stain electron microscopy **(A)** and bright-field confocal microscopy **(B)** recordings of *E. coli* WT-like, locked-ON and locked-OFF cells. A white 500□nm **(A)** and a black 2 μm **(B)** scale bar is included in each image. **(C)** Quantitative analyses of cells displaying fimbriae at their surface for the WT-like (n = 54), locked-ON (n = 54) and locked-OFF (n=56) strains. (**D**) Yeast agglutination assay with WT-like, locked-ON and locked-OFF *E. coli* cells in the absence or presence of 5% of the FimH competitive ligand D-mannose. (**E**) Growth curves of WT-like (black dot), Δ*fimE* (grey diamond), locked-ON (blue triangle) and locked-OFF (red inversed triangle) *E. coli* strains. The data represent the mean (±SD) of three independent experiments.

### Fimbriae-mediated microcolonies resistance to various T6SS

To further analyze the capacity of type 1 fimbriae-mediated microcolonies to confer resistance against T6SS-mediated attacks, and to exclude that they specifically resist to the set of effectors of CM3267, we performed interbacterial competition assays using the WT-like, Δ*fimE,* locked-ON and locked-OFF strains as targets, and the T6SS⁺ attackers CM3267, EAEC 17-2, and *C. rodentium*. Target survival was measured using the Survival Growth Kinetics (SGK) assay, a method based on the time of emergence of the target after coincubation with the attacker, which is inversely proportional to the number of target cells that have survived the attacks ^16^.

Competition assays showed that the WT-like *E. coli* target strain is efficiently killed by all the attackers in a T6SS-dependent manner (Figure 2A, C, E). In all cases, strains producing type 1 fimbriae (Δ*fimE* and locked-ON) consistently showed higher survival than low- or non-fimbriated strains (WT-like and locked-OFF). Because the three attackers exhibited different killing efficiencies against *E. coli*, we calculated the relative resistance by normalizing the survival of the locked-OFF, locked-ON and Δ*fimE* strains to that of the WT-like target. In all competition assays, the locked-OFF strain displayed susceptibility comparable to the WT-like strain, whereas the locked-ON and Δ*fimE* strains survive 1.5-2-fold better (Figure 2B, D, F). Collectively, these results indicate that type 1 fimbriae-mediated microcolony formation confers increased protection against T6SS-mediated attacks from various species, independently of their sets of effectors.

**Figure 2.**
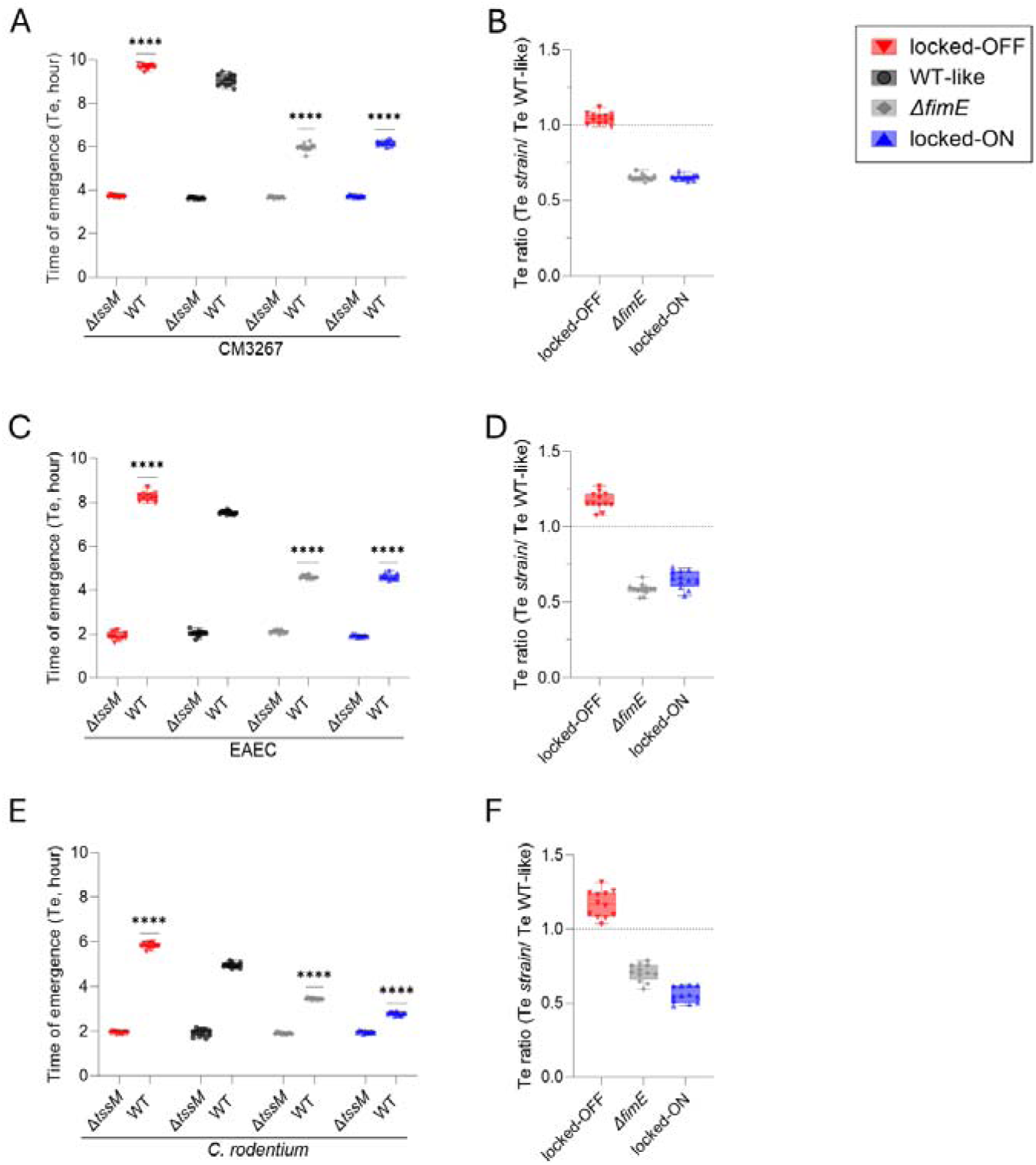
Type 1 fimbriae confer protection against T6SS mediated attacks. Interbacterial competition assays **(A-C-E)** and normalized survival ratios (target/WT-like) **(B-D-F)** of WT-like (black), Δ*fimE* (gray), locked-ON (blue) or locked-OFF (red) target cells after co-incubation with CM3267 **(A-B)**, EAEC 17-2 **(C-D)** or *C. rodentium* RLC55 **(E-F)** attackers. Box-and-whisker representation of the target cell time of emergence (Te, hours) (**A**) and Te ratio (**B**) (horizontal bar, median value; lower and upper boundaries of the box plot, 25% and 75% percentiles, respectively; whiskers, 10% and 90% percentiles). The independent raw values from four independent experiments (3 replicates for each) are shown as plain circles, triangles or diamonds. Statistical significance was assessed using a two-way (A, C, E) ANOVA test (*\*\*\**, p< 0.001; *\*\*\*\**, p< 0.0001). Comparisons were made against WT-like controls (A, C, E).

### Microcolonies shield against other contact-dependent weapons

To test whether microcolonies confer protection beyond T6SS attacks, we next performed competition assays between our target strains and attackers using other contact-dependent antibacterial mechanisms: *Lysobacter enzymogenes* OH11, a soilborne biocontrol bacterium equipped with a functional antibacterial T4SS ^17,18^; and *E. coli* producing a contact-dependent growth inhibition (CDI) system composed of the CdiB transporter and a chimera CdiA passenger exposing the antibacterial *Xenorhabdus bovienii* TreX ADP-ribosyltransferase that targets FtsZ ^19,20^. Figure 3 shows that non-fimbriated target cells (WT-like and locked-OFF) were susceptible to T4SS (Figure 3A-B) and CDI (Figure 3C-D) attacks, whereas fimbriated Δ*fimE* and locked-ON strains exhibited higher survival. In both cases, the locked-OFF strain mirrored the susceptibility of the WT-like strain when survival ratios were normalized to the WT-like control. In contrast, both the locked-ON and Δ*fimE* target strains showed enhanced survival. It is worth to note that the locked-ON strain displayed greater protection than the Δ*fimE* strain, indicating that cells organized in more robust microcolonies are better shielded against T4SS- and CDI-mediated attacks (Figure 3B-D). Overall, these findings indicate that, similarly to the resistance conferred to T6SS attacks, microcolonies protect target bacteria from other contact-dependent competition mechanisms, including T4SS and CDI.

**Figure 3.**
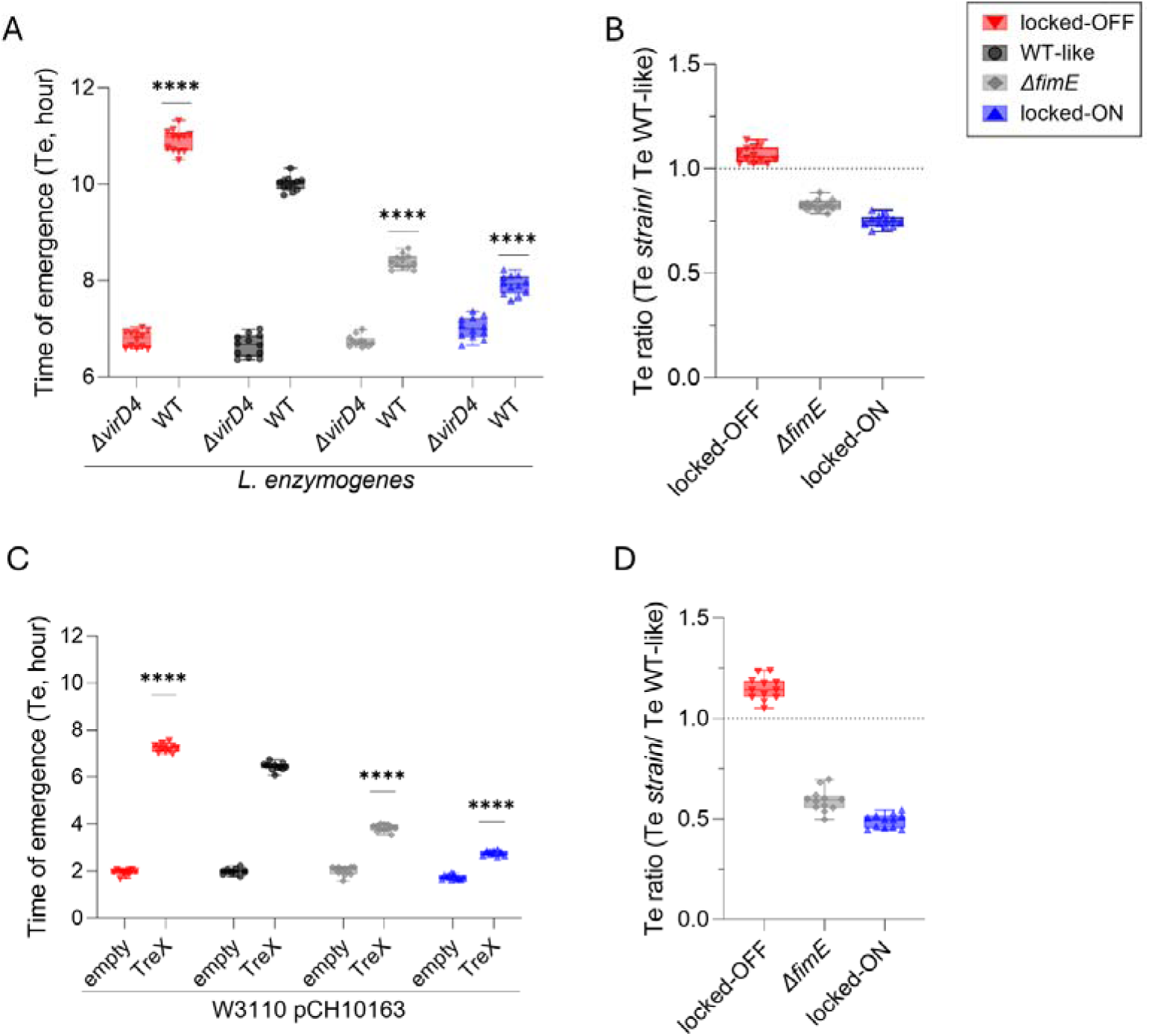
Type 1 fimbriae confer protection against short-range weapons such as CDI and T4SS. Interbacterial competition assay **(A, C)** and normalized survival ratios **(B, D)** of WT-like (black), Δ*fimE* (gray), locked-ON (blue) or locked-OFF (red) target cells after co-incubation with *Lysobacter enzymogenes* OH11 **(A)** or *E. coli* W3110 pCH10163-TreX **(C)** attackers. Box-and-whisker representation of the target cell time of emergence (Te, hours) **(A, C)** and Te ratio **(B, D)** (horizontal bar, median value; lower and upper boundaries of the box plot, 25% and 75% percentiles, respectively; whiskers, 10% and 90% percentiles). The independent raw values from four independent experiments (3 replicates for each) are shown as plain circles, triangles, or diamonds. Statistical significance was assessed using a two-way (A, C) ANOVA test (****, p< 0.0001). Comparisons were made against WT-like controls (A, C).

### Microcolonies provide limited protection against soluble toxins

Bacterial antagonism is not limited to contact-dependent strategies but can also involve diffusible toxins or compounds, such as colicins, toxins secreted by certain *E. coli* strains that are lethal to closely related strains ^21^, and antibiotics. Previous studies have shown that colicins can disrupt established bacterial biofilm ^22^ and kill sensitive cells within mixed-species biofilms ^23^. On the other hand, biofilms prevent antibiotics from reaching bacterial cells, and hence correlate with increased antibiotic tolerance ^10,24^. Based on these observations, we next sought to determine whether colicins and antibiotics could eliminate bacterial cells embedded within microcolonies. Survival of target WT-like, locked-OFF, locked-ON, Δ*fimE* target cells was measured after exposure to different colicins or antibiotics. Upon incubation with different classes of colicins (colicin A, Tol-dependent pore-forming activity; colicin E9, Tol-dependent nuclease activity; colicin M, TonB-dependent lipid A hydrolase) at a cell:colicin ratio of 1:1000 for 30 minutes, survival was low for all target cells and no differences in killing efficiency were detected (Figure 4A-B).

**Figure 4.**
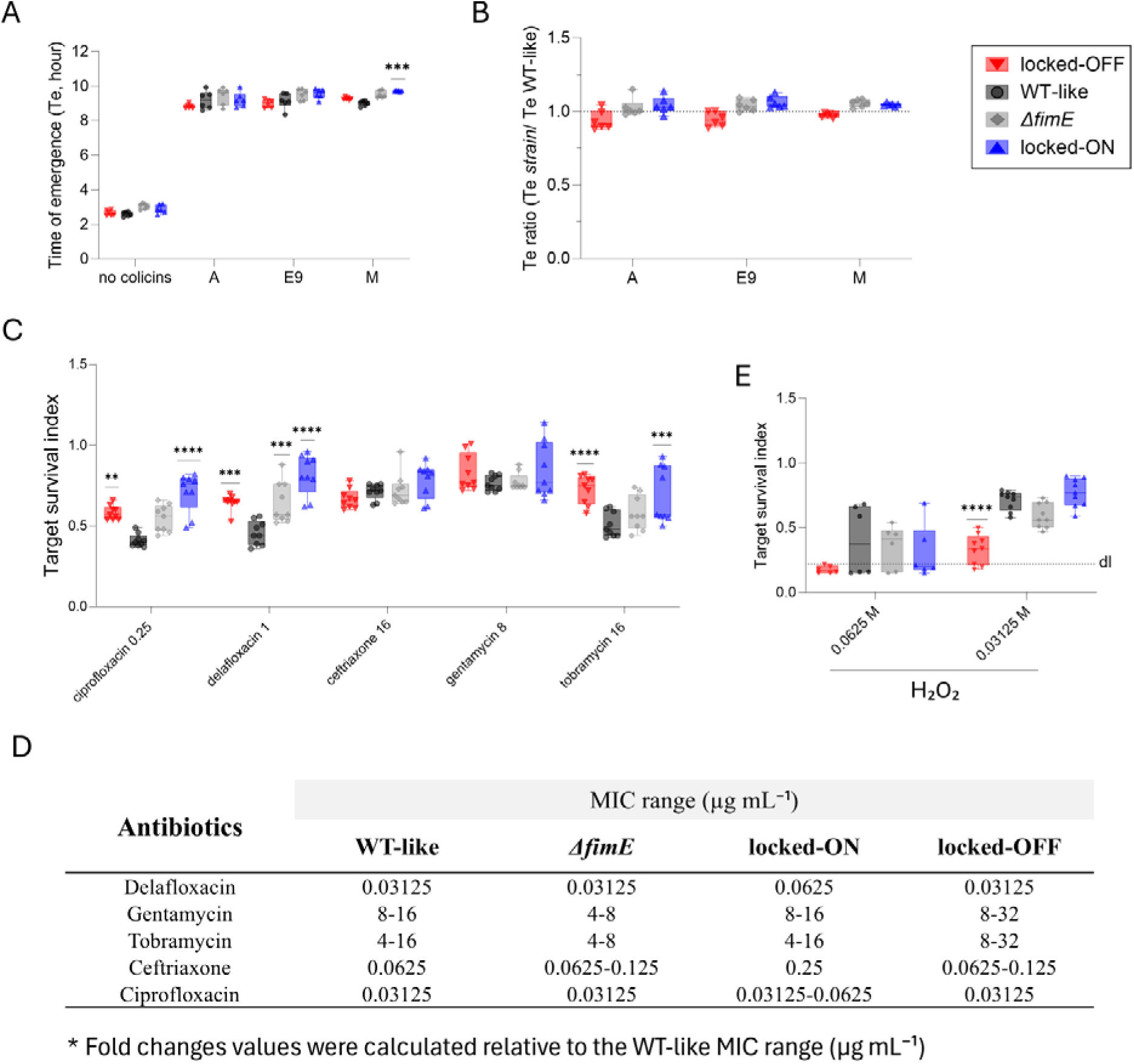
Long-range diffusible toxins affect the susceptibility of cells embedded within microcolonies. **(A)** Survival of WT-like (black), Δ*fimE* (gray), locked-ON (blue), and locked-OFF (red) target cells following exposure to colicins A, E9, and M. **(B)** Corresponding normalized survival ratios for the same target strains. **(C)** Target survival index of WT-like (black), Δ*fimE* (gray), locked-ON (blue), and locked-OFF (red) cells after 90 min exposure to ciprofloxacin (0.25 µg mL^-^¹), delafloxacin (1 µg mL^-^¹), ceftriaxone (16 µg mL^-^¹), gentamicin (8 µg mL^-^¹), and tobramycin (16 µg mL^-^¹). **(D)** Table summarizing the MIC range of the antibiotics against WT-like, Δ*fimE*, locked-ON, and locked-OFF strains. **(E)** Target survival index of WT-like (black), Δ*fimE* (gray), locked-ON (blue), and locked-OFF (red) cells after exposure to H_₂_O_₂_ (0.03125 M and 0.0625 M).Box-and-whisker representation, horizontal bar, median value; lower and upper boundaries of the box plot, 25% and 75% percentiles, respectively; whiskers, 10% and 90% percentiles. The independent raw values from six independent experiments are shown as plain triangles, circles, or diamonds. Statistical significance was assessed using a two-way **(A, C, E)** ANOVA test (**, p< 0.01; ***, p< 0.001; *\*\*\*\**, p< 0.0001). Comparisons were made against WT-like controls **(A, C, E)**.

Similarly, fimbriae-mediated microcolony formation failed to provide significant protection upon exposure to different classes of antibiotics, including ciprofloxacin and delafloxacin fluoroquinolones targeting DNA replication, tobramycin and gentamycin aminoglycosides inhibiting protein synthesis, and ceftriaxone a third generation of cephalosporin antibiotic preventing cell wall synthesis (Figure 4C). While the survival index of each target strain varied depending on the antibiotic tested, we did observe a small increase in short-term survival for ciprofloxacin, delafloxacin and tobramycin. For these antibiotics, Δ*fimE,* and locked-ON strains showed higher short-term survival than the WT-like strain, with the locked-ON strain generally exhibiting slightly higher survival than the Δ*fimE* mutant. Surprisingly, the locked-OFF mutants also showed higher survival than the WT-like bacteria. In contrast, no significant differences in survival were observed among the four strains when exposed to ceftriaxone or gentamicin. To determine whether this trend persisted under longer-term exposure, we performed MIC assays. Overall, MIC values were comparable for all strains, with differences generally within ∼2-fold, which is typically within the standard error for these assays. The one exception was ceftriaxone, for which the locked-ON strain exhibited a 4-fold higher MIC value than the WT-like strain. However, this increase is specific to ceftriaxone and does not appear to reflect a general resistance mechanism conferred by microcolony formation.

Finally, an oxidative stress survival assay showed that the locked-OFF strain exhibited lower survival than WT-like, Δ*fimE*, and locked-ON strains at both 0.03125 M and 0.0625 M of H₂O₂. No significant differences were observed among WT-like, Δ*fimE*, and locked-ON strains under these conditions, indicating that type 1 fimbriae-mediated microcolonies do not confer substantial protection against oxidative stress.

Collectively, these results indicate that diffusible toxic compounds and proteins are capable of penetrating type 1 fimbriae-mediated microcolonies and of efficiently killing aggregated bacteria.

### Heterogeneity of type 1 fimbriae expression in microcolonies

Due to phase variation controlled by the FimB and FimE recombinases and by additional elements, clonal populations of *E. coli* are heterogeneous for *fim* operon expression ^14,15^. We therefore wondered whether *fim* heterogeneity exists within microcolonies, and if so, whether non-fimbriated cells were sheltered inside to benefit from the collective protection conferred by fimbriated cells.

To monitor *fim* heterogeneity, we engineered a reporter vector, pColorSwitch, carrying genes encoding the mRFP and sfGFP fluorescent proteins on both sides of the *fimS* promoter (Figure 5A). In this construct, the promoter expresses the mRFP gene when in the OFF position and the sfGFP gene when switched to the ON position. In agreement with previous observations, confocal microscopy confirmed that WT-like cells bearing the pColorSwitch vector were mostly in the OFF state and did not aggregate (Figure 5B-C). In contrast, Δ*fimE* cells carrying pColorSwitch were more heterogeneous and formed microcolonies containing both sfGFP⁺ and mRFP⁺ cells (Figures 5B-C), indicating that bacteria in the *fim* OFF state are sheltered within microcolonies stabilized by bacteria in the *fim* ON state. It is worth to note that cells producing both mRFP and sfGFP can be observed in the WT-like and Δ*fimE* background (Figure 5B-C). Moreover, the proportions of sfGFP⁺, mRFP⁺, or dual sfGFP⁺/mRFP⁺ cells observed at the single-cell level matched those obtained from colony fluorescence analysis on agar plates (Figures 5C-D). Collectively, these results support the idea that type 1 fimbriae-mediated microcolonies promote phenotypic heterogeneity within bacterial communities.

**Figure 5.**
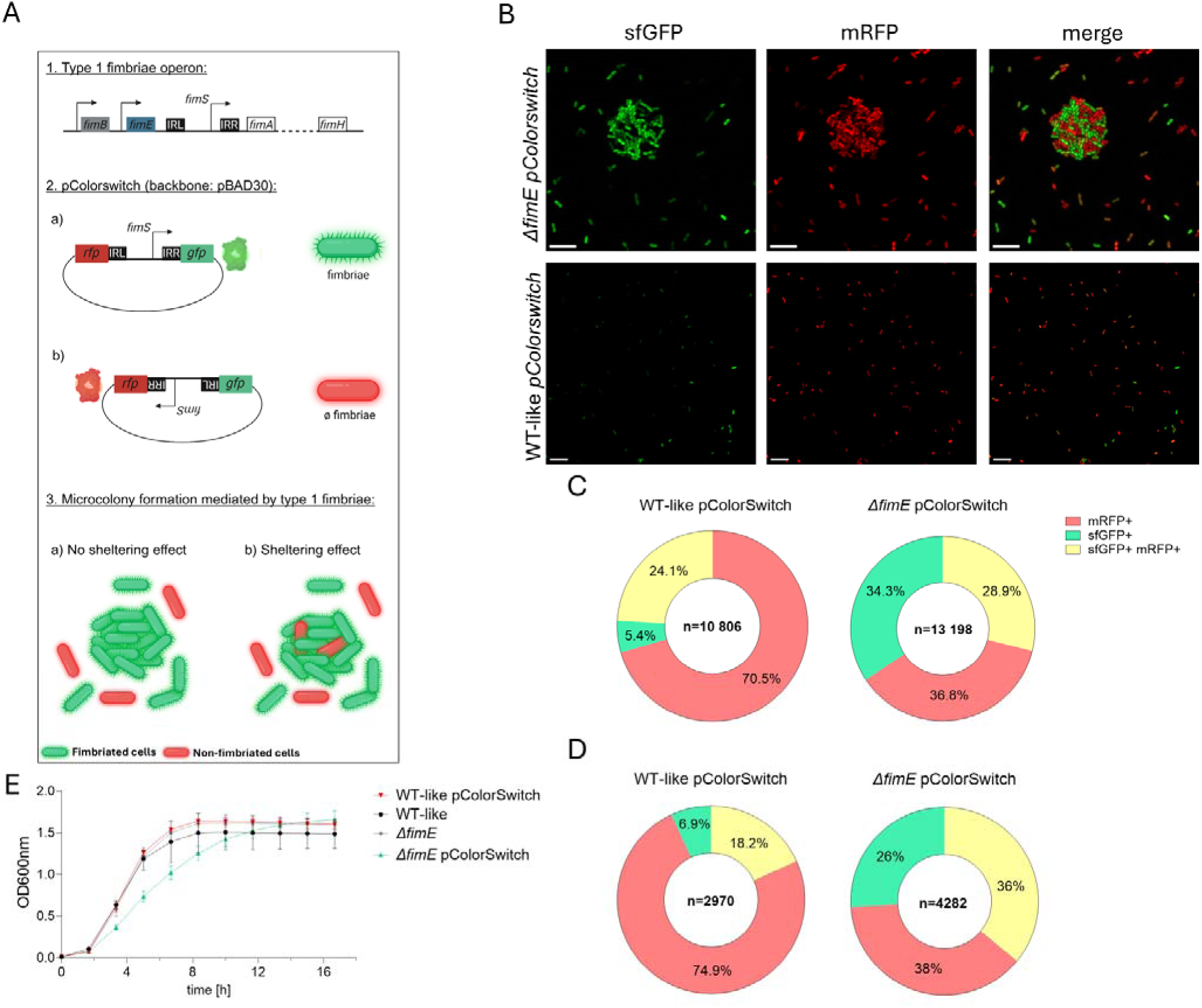
Type 1 fimbriae-mediated microcolonies are heterogeneity supporting environments. **(A)** The *fimS* promoter was amplified and cloned into a pBAD30 backbone carrying mRFP and sfGFP fluorescent proteins on both side of the *fimS* promoter (1). If the promoter is turned in the ON orientation, sfGFP is produced (2a). If the promoter is turned in the OFF orientation, mRFP is produced (2b). This construct allows monitoring of phase variation in *E. coli* BW25113 microcolonies (3) We used this to test if non type 1 fimbriae-producing bacteria could be embedded within microcolonies, profiting a sheltering effect or not. **(B)** Overnight cultures of WT-like and Δ*fimE* strains carrying the pColorSwitch reporter were centrifuged, vortexed, spotted in triplicate, and observed by epifluorescence microscopy. Each images includes a white 5□μm scale bar. **(C)** Proportion of sfGFP⁺, mRFP⁺, and dual sfGFP⁺/mRFP⁺ cells in WT-like (n = 10,806) and Δ*fimE* (n = 13,198) cells carrying the pColorSwitch reporter. **(D)** Proportion of sfGFP⁺, mRFP⁺, and dual sfGFP⁺/mRFP⁺ colonies on agar plates in WT-like (n = 2,970) and Δ*fimE* (n = 4,282) strains carrying pColorSwitch. **(E)** Growth curves of WT-like and Δ*fimE* cells with or without the pColorSwitch reporter is shown. Overnight cultures were diluted 1:100, transferred to 96-well plates, and grown for 17 h. OD₆₀₀ was measured every 20 min. Data represent the mean ± SD of three independent experiments.

### Fim^-^ cells are protected from T6SS within microcolonies

To further test whether sheltered Fim^-^ cells can benefit from the protection conferred by the microcolony, we compared the survival of locked-OFF cells alone or mixed in a 1:1 ratio with the locked-ON cells when exposed to CM3267 T6SS attacks. To distinguish both strains, locked-ON and locked-OFF cells were transformed with distinct fluorescent plasmids. As previously observed, locked-ON fluorescent cells exhibited a high target survival index (0.42), while locked-OFF fluorescent cells had low survival (0.1) when tested alone against CM3267. However, in mixed cultures, locked-OFF fluorescent cells displayed significantly improved survival (0.24) (Figure 6A-C).

**Figure 6.**
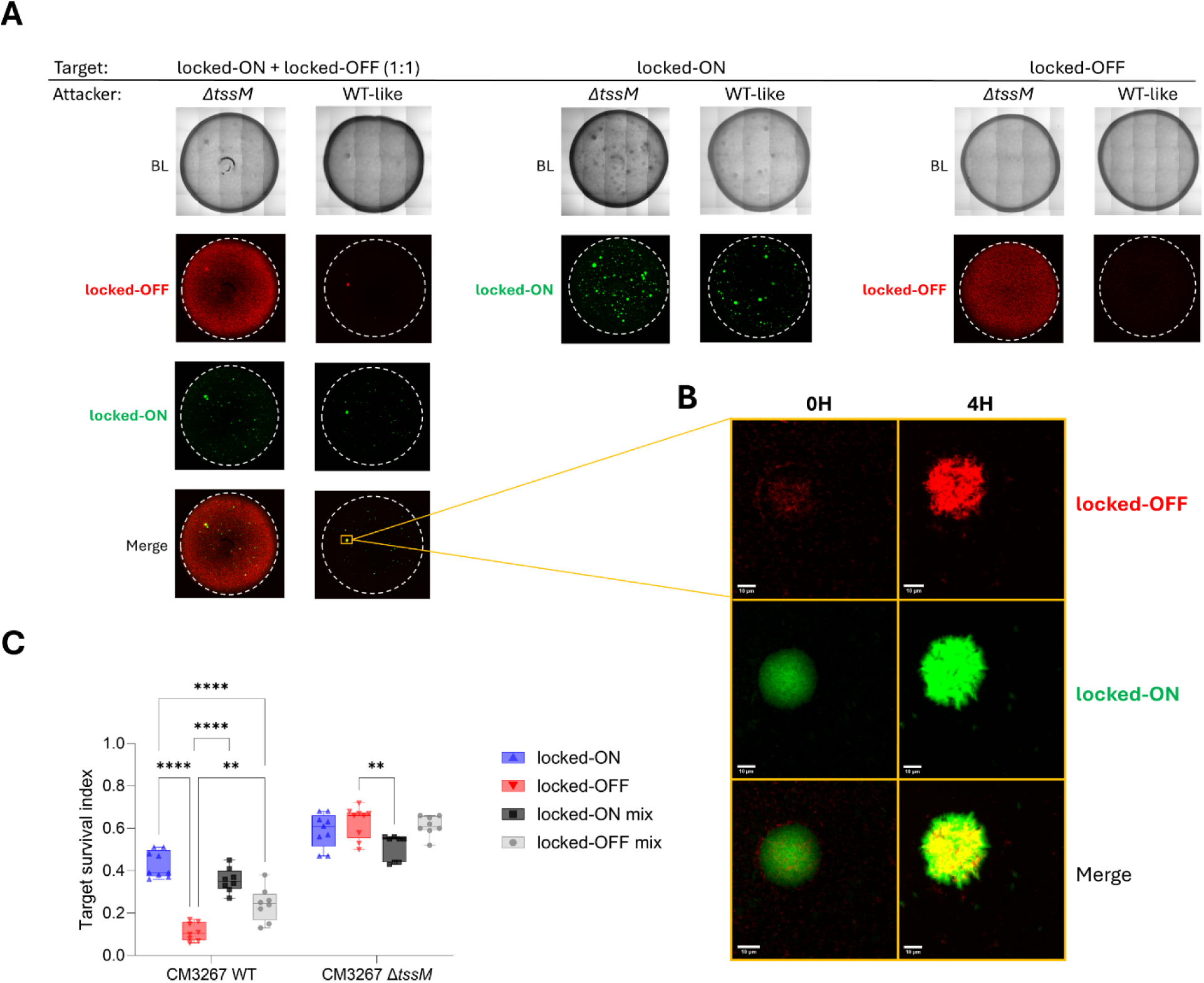
Lock-OFF cells sheltered inside microcolonies formed by type 1 fimbriae producing *E. coli* cells are protected from T6SS attacks. **(A)** Microscopy images of competition spots of fluorescently labeled locked-OFF (red), locked-ON (green) or mixed locked-ON/locked-OFF target cells in the presence of Δ*tssM* or WT CM3267 attackers at times=0 and 4 hours. **(B)** 2× close-up of the locked-ON/locked-OFF colony at 0 and 4 h post competition with the CM3267 WT attacker. Each zoomed images includes a white 10□μm scale bar. **(C)** Interbacterial competition assay between CM3267 attacker and fluorescently labeled locked-ON (green triangle), locked-OFF (red inverted triangle) or mixed locked-ON/locked-OFF target cells. Box-and-whisker representation of the target survival index (horizontal bar, median value; lower and upper boundaries of the box plot, 25% and 75% percentiles, respectively; whiskers, 10% and 90% percentiles). The independent raw values from three independent experiments (3 replicates for each) are shown as plain triangles, squares and circles. Statistical significance was assessed using a two-way ANOVA test (ns, non-significant; **, p< 0.01; *\*\*\*\**, p< 0.0001).

To gain further evidence for locked-OFF protection within microcolonies, we examined competition spots by confocal microscopy (Figure 6A). Against CM3267 Δ*tssM* attackers, locked-ON cells appeared both as microcolonies and single cells, whereas lock-OFF cells remained only as isolated bacteria. Against wild-type CM3267, single cells of the locked-ON strain were eliminated and only microcolonies persisted, while locked-OFF bacteria were almost completely eradicated (Figure 6A). Finally, in mixed cultures, both fluorescence signals colocalized within microcolonies. Magnification of microcolonies showed that locked-OFF cells that were trapped inside locked-ON aggregates were protected during the 4 hours of the competition assay (Figure 6B).

Together, these results demonstrate that fimbriated cells form microcolonies that resist *C. malonaticus* T6SS attacks but also shelter non-fimbriated cells, protecting them and supporting their growth. This cooperative protection enables the persistence and propagation of phenotypic diversity within bacterial populations under competitive conditions.

### Aggregation-promoting structures mediates resistance to T6SS

Other surface structures such as autotransporters can also promote the aggregation of cells. Our study thus also raises the question of whether other aggregation-promoting structures also confer resistance to T6SS. We therefore investigated if TibA and Antigen 43 (Ag43), two T5SS autotransporter adhesins that promote self-aggregation ^25–27^, could also contribute to bacterial resistance against T6SS-mediated assaults. As expected, we observed using confocal microscopy that WT-like cells producing the TibA or Ag43 adhesins promoted aggregation (Figure 7A). We then performed interbacterial competition assays between the *E. coli* strains producing, or not, TibA or Ag43, as targets and CM3267 as attacker. Figure 7C shows that despite a slower growth (Figure 7B), cells producing the TibA or Ag43 adhesins were more resistant to CM3267 T6SS attacks compared to cells carrying the empty vector (Figure 7C). Collectively, these results show that, similar to type 1 fimbriae, the TibA and Ag43 adhesins promote microcolony formation and confer resistance to T6SS-mediated attacks from CM3267.

**Figure 7.**
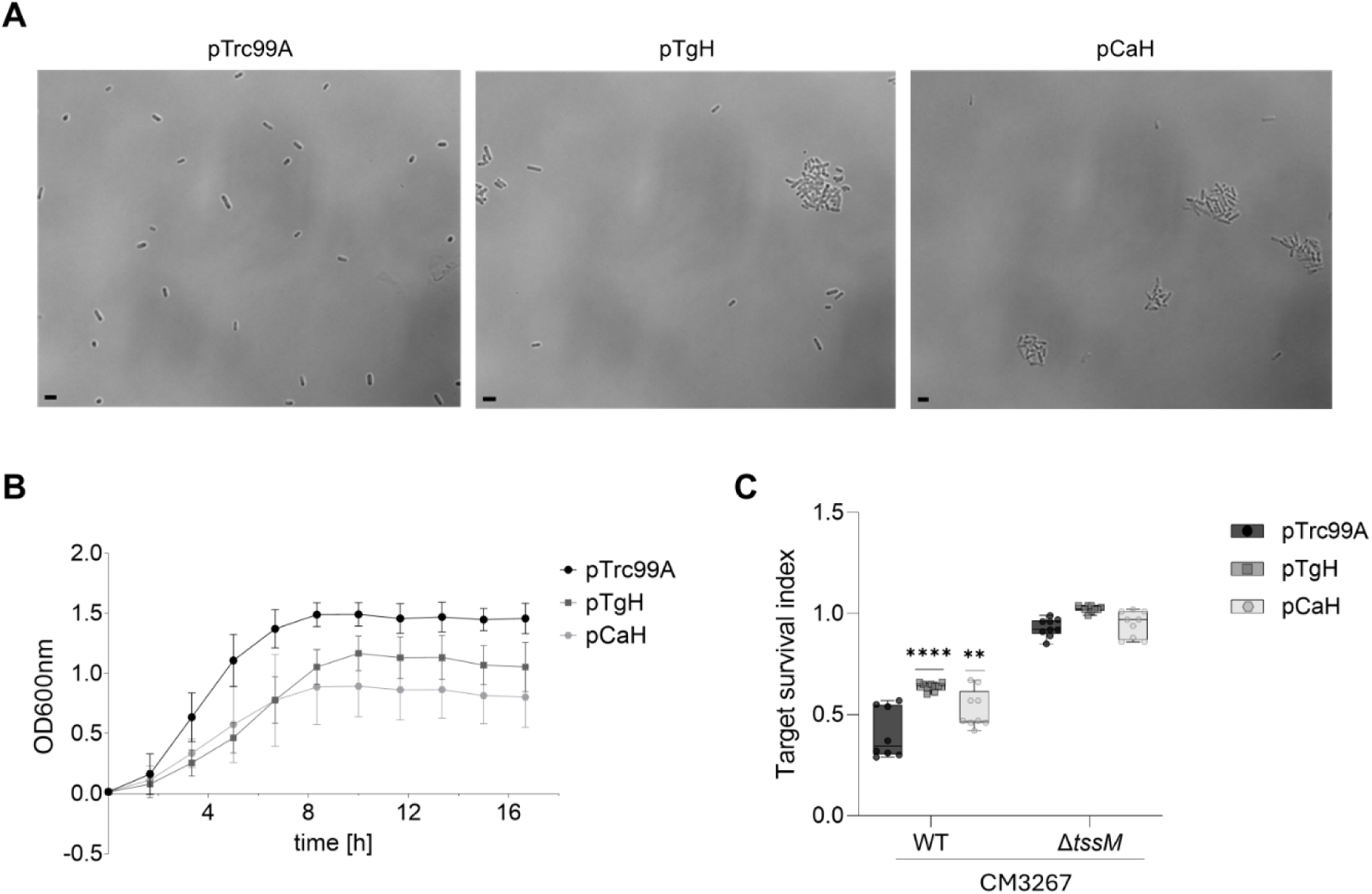
The TibA (pTgH) and Ag43 (pCah) T5SS adhesins confer resistance against C. malonaticus 3267 T6SS attacks. (**A**) Representative bright-field confocal microscopy recordings of WT-like *E. coli* target cells carrying the empty pTrc99A vector (empty) or pTrc99A derivatives pTgH (TibA) or pCaH (Ag43). Each images includes a black 2□μm scale bar. (**B**) Growth curves of WT-like *E. coli* strains carrying the pTrc99A vector (empty) (black dot) or pTrc99A derivatives pTgH (TibA) (dark grey square) or pCaH (Ag43) (light grey square). The data represent the mean (±s.d.) of three independent experiments. (**C**) Interbacterial competition assay between CM3267 WT or Δ*tssM* attackers and WT-like *E. coli* target cells carrying the pTrc99A vector (empty) or pTrc99A derivatives producing TibA (pTgH) or Ag43 (pCaH). Box-and-whisker representation of the target survival index (horizontal bar, median value; lower and upper boundaries of the box plot, 25% and 75% percentiles, respectively; whiskers, 10% and 90% percentiles). The independent raw values from three independent experiments (3 replicates for each) are shown as plain squares, circles, or diamonds. Statistical significance was assessed using a two-way ANOVA test (**, p< 0.01; ****, p< 0.0001).

### Aggregation phenotypes vary among clinical bacterial strains

T6SS-mediated antagonism is a key strategy employed by the gut microbiome to limit the colonization by pathogens ^28,29^. We thus wondered whether different pathogenic strains could aggregate and form microcolonies. We visualized the aggregation of 12 different species with varying clinical outcomes and various hosts, such as avian, porcine, and human hosts, and observed different levels of aggregation (Figure 8A). In particular, *E. coli* 17-2 and LF82, respectively EAEC and AIEC pathotypes, and *Acinetobacter baumannii* formed large aggregates. Other strains formed smaller aggregates (*E. coli* 2787, *Citrobacter rodentium* DSB100 or *Klebsiella oxytoca* for instance) and there were a few strains for which we did not observe aggregation (*E. coli* H10407, *E. coli* PD20 for instance). Overall, most clinical strains formed aggregates, suggesting that aggregation might represent a wide-spread mechanism to resist contact-dependant antagonism.

**Figure 8.**
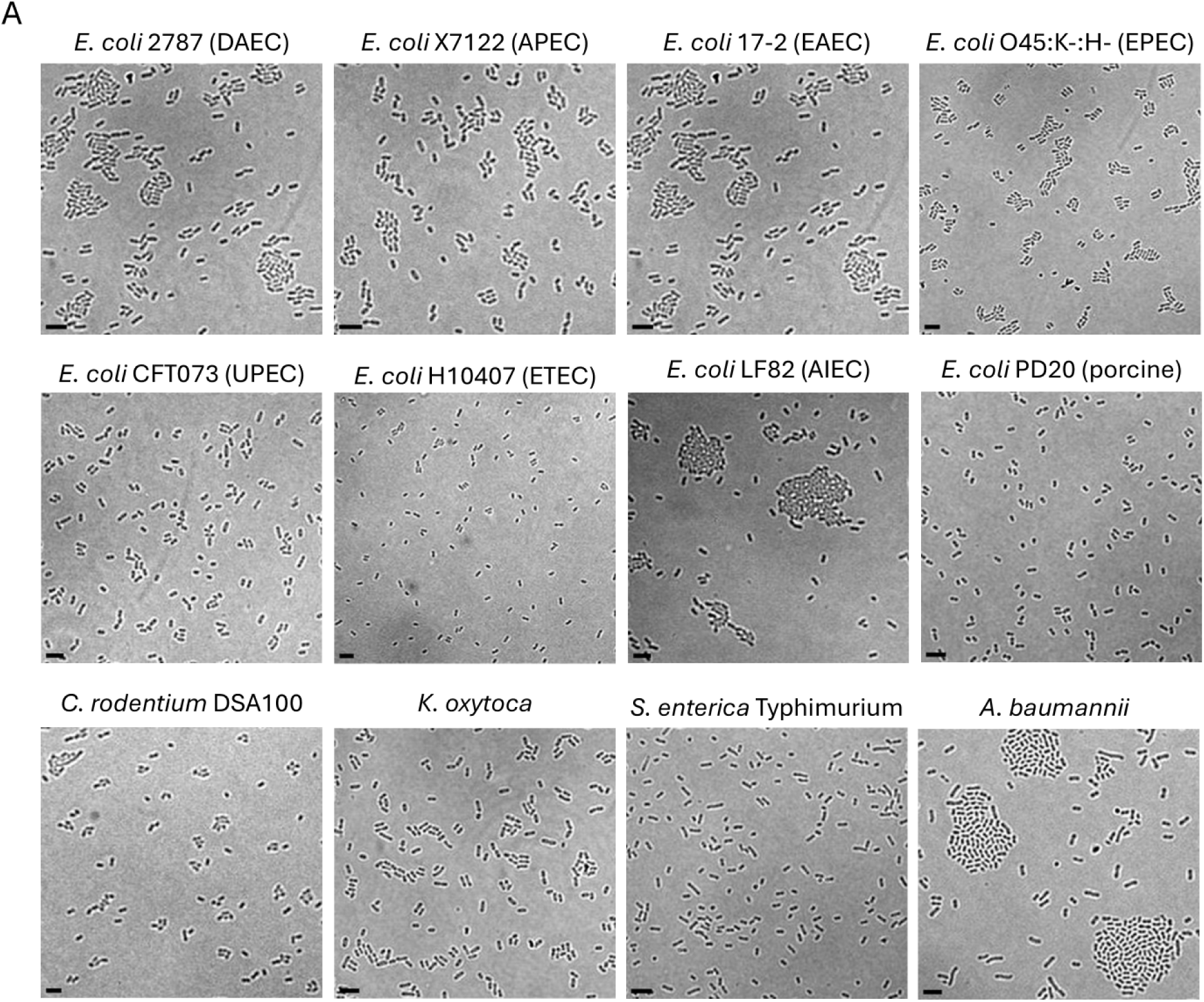
Species- and strain-dependent aggregation among pathogenic bacteria. Representative bright-field confocal microscopy images of overnight cultures of *Escherichia coli* strains 2787, X7122, 17-2, O45:K−:H−, CFT073, H10407, LF82, and PD20, *Citrobacter rodentium* DSA100, *Klebsiella oxytoca*, *Salmonella enterica* serovar Typhimurium, and *Acinetobacter baumannii*. Each images includes a black 2□μm scale bar.

## Discussion

Deciphering how and why bacteria adhere together is an important parameter to understand the dynamics of microbial communities. Specifically, biofilms that emerge from prior adhesion to surface or bacterial aggregation are generally critical for bacterial survival and pathogenesis, and shape cooperative and antagonistic behaviors in communities. For instance, we have previously identified that a *fimE* deletion mutant overexpressing type 1 fimbriae forms microcolonies that confer collective protection from *Cronobacter malonaticus* T6SS assaults ^13^.

Here, we extend these observations to demonstrate that type 1 fimbriae–mediated microcolonies provide resistance against the T6SS from different species (Figure 2), which deploy distinct effector repertoires: phospholipases in EAEC ^30^, amidases and metallopeptidases in *C. rodentium* ^31^, and likely amidases and phospholipases in *C. malonaticus*. This argues against resistance to a specific toxic activity but instead supports a mechanical mode of protection against T6SS attacks. Interestingly, microcolonies protection was not limited to T6SS attacks. Indeed, we also observed that aggregation led to target survival against other toxin delivery systems, such as T4SS and CDI, which also rely on direct cell-cell contact for efficient intoxication of target cells (Figure 3). Aggregation can also protect target cells against microbial predation ^32^. In contrast, microcolonies failed to protect cells against diffusible toxins such as colicins. Colicins are antibacterial toxins released by producing cells, which display enzymatic (colicin E9 – DNase^21^ and colicin M - PG precursors degradation^33^) or pore-forming (colicin A^34,35^) activities. We observed that aggregated bacteria did not survive exposure to these diffusible toxins, regardless of the targeted cellular compartment. Similarly, other soluble molecules such as H₂O₂ or antibiotics also disrupted microcolonies (Figure 4).

In the case of antibiotics, we observed a small gain in survival with the *fimE* mutant or locked-ON cells for some antibiotics, but these effects were transient as MIC assays showed no significant differences. This is in contrast to aggregated *Staphylococcus aureus* cells, which exhibit up to a 30-fold increased survival when exposed to ciprofloxacin compared to non-aggregated cells ^10^. This discrepancy could be due to differences in the composition of the microcolony, such as the presence of EPS and to the much larger number of cells typically involved in these aggregates. Interestingly, the locked-OFF strain also survived better in the short-term than the WT-like strain against some antibiotics. Locked-OFF cells were also extremely sensitive to H₂O₂. Locked-OFF cells did not exhibit any growth defect by themselves. Thus, this increased sensitivity to H₂O₂ could suggest a possible link between the metabolic state of the cells and an increased susceptibility to ROS induced by H₂O₂.

During our experiments, we used a panel of target cells that displayed a gradient of fimbriae expression, from locked-OFF cells (no fimbriae), WT-like cells (∼5% of fimbriated cells; low density of fimbriae/cell), Δ*fimE* (∼50% of fimbriated cells; low to high density of fimbriae/cell) and locked-ON cells (100% fimbriated cells; high density of fimbriae/cell). In general, resistance against contact-dependent attacks correlated with fimbriae expression. In a few cases (EAEC, *C. rodentium*), the Δ*fimE* and locked-ON strains displayed comparable levels of resistance. However, locked-ON cells exhibited a slight growth defect, which may account for this observation. This slight growth defect suggests a fitness cost associated with high-level fimbriae production, consistent with a trade-off between microcolony-mediated protection and the metabolic burden of type 1 fimbriae expression. Such microcolonies may therefore promote phenotypic heterogeneity, allowing non-fimbriated cells to be sheltered within the community while benefiting from collective protection.

Cheater bacteria have been documented not only in laboratory systems ^36,37^ but also in natural communities. Recently, Butaitè *et al.* ^38^ showed that in soil and pond communities of *Pseudomonas* species, some isolates produce little or no pyoverdine, an iron scavenging compound, yet still express specific pyoverdine receptors. In these communities, where some *Pseudomonas* isolates produce large amount of pyoverdine, isolates that produce no or low levels of pyoverdine can act as cheaters, using this shared public good to survive ^38^.

To get information on the sheltering effect, we engineered a pColorSwitch system for real-time monitoring of *fimS* switching events, which allowed to observe cells with the promoter in the ON- and OFF-states within the microcolonies (Figure 5). Importantly, the proportion of *fimS* ON and OFF cells observed by single-cell fluorescence matched the switching frequencies measured at the colony-phenotype level validating that the single-cell pColorSwitch readout reflects the population-level *fimS* state. However, we also observed cells producing both mRFP and sfGFP in the WT-like and Δ*fimE* backgrounds, likely due to the long half-lives of mRFP and sfGFP and/or the presence pColorSwitch copies carrying the *fimS* promoter in different orientations within the same cell.

To further observe if non-fimbriated cells could be sheltered inside microcolonies, we mixed locked-ON and locked-OFF cells (1:1 ratio) (Figure 6). Again, we did observe that locked-OFF cells were part of the microcolonies formed by the locked-ON cells. Moreover, locked-OFF cells embedded within the microcolonies were protected against T6SS attacks. Furthermore, locked-ON survival was unchanged whether they were challenged by the T6SS alone or in a 1: 1 mixture with locked-OFF strain, indicating that the presence of sheltered bacteria did not decrease the overall protection provided by microcolonies. These results suggest that only a fraction of fimbriae-producing cells is required for the population to resist T6SS attacks. However, the stability of this protective effect under different population compositions (i.e. increasing proportions of locked-OFF cells) remains unclear. Similar thresholds have been described in other cooperative systems. For example, EPS-mediated pellicle formation at the air-liquid interface depends on a sufficient proportion of EPS-producing cells. Cheater bacteria that do not produce EPS but benefit from it and often exhibit higher fitness can persist within the pellicle, but when their proportion exceeds a critical threshold, the structure collapses and the collective benefit of EPS production is lost because the producer minority cannot sustain the biomass ^39,40^. Additional experiments are needed to address this question. Notably, the locked-ON strain exhibits a survival defect in the mixed condition locked-ON/locked-OFF against the T6SS-defective attacker, indicating that microcolony-mediated protection arises only during competition with a T6SS⁺ attacker. In the absence of such competition, the locked-ON strain seems disadvantaged, consistent with the previously observed trade-off.

Interestingly, other structures promoting aggregation also confer resistance to T6SS. Here, we observed that T5SS adhesins, known to promote strong self-aggregation, also provided resistance against T6SS assaults from CM3267 (Figure 7). Similarly, *Vibrio cholerae* can use their type 4 pili to survive T6SS attacks through spatial segregation ^41^. This suggests that structures that promote aggregation are capable of conferring resistance to T6SS and other contact-dependent antibacterial systems. In nature, aggregation is a widely observed phenomenon, that can be mediated by a large range of adhesive structures ^42^. We additionally observed aggregation of a range of pathogenic bacteria such as EAEC, ETEC, EHEC and *Acinetobacter baumannii*, with some strains forming large aggregates ^43,44^. While *E. coli* H10407 and PD20 did not aggregate in our assay, they are known to produce TibA and AIDA-I respectively, which are both self-associating autotransporters. The expression level of TibA and AIDA-I in these strains in laboratory conditions such as those tested here is not known. However, when TibA and AIDA-I are expressed, both strains form large aggregate ^45^. Overall, this suggests that cell aggregation is potentially found in host-associated environments and could represent a resistance mechanism against competition mechanisms (T6SS, T4SS, CDI) that usually occur in dense microbial population ^28,46,47^. Consistent with this hypothesis, a recent study has shown that bacteria can sense signals from potential attacker bacteria and react to this signal by forming aggregation to increase their survival ^48^.

Finally, our work highlights the protective and social functions of microcolonies, providing new insights into bacterial survival strategies under competitive and hostile conditions. These structures not only safeguard bacteria from contact-dependent antagonists but can also promote phenotypic diversity within microbial communities (Figure 9). Deciphering microcolony-mediated resistance has important implications for understanding dynamic interaction in polymicrobial communities.

**Figure 9.**
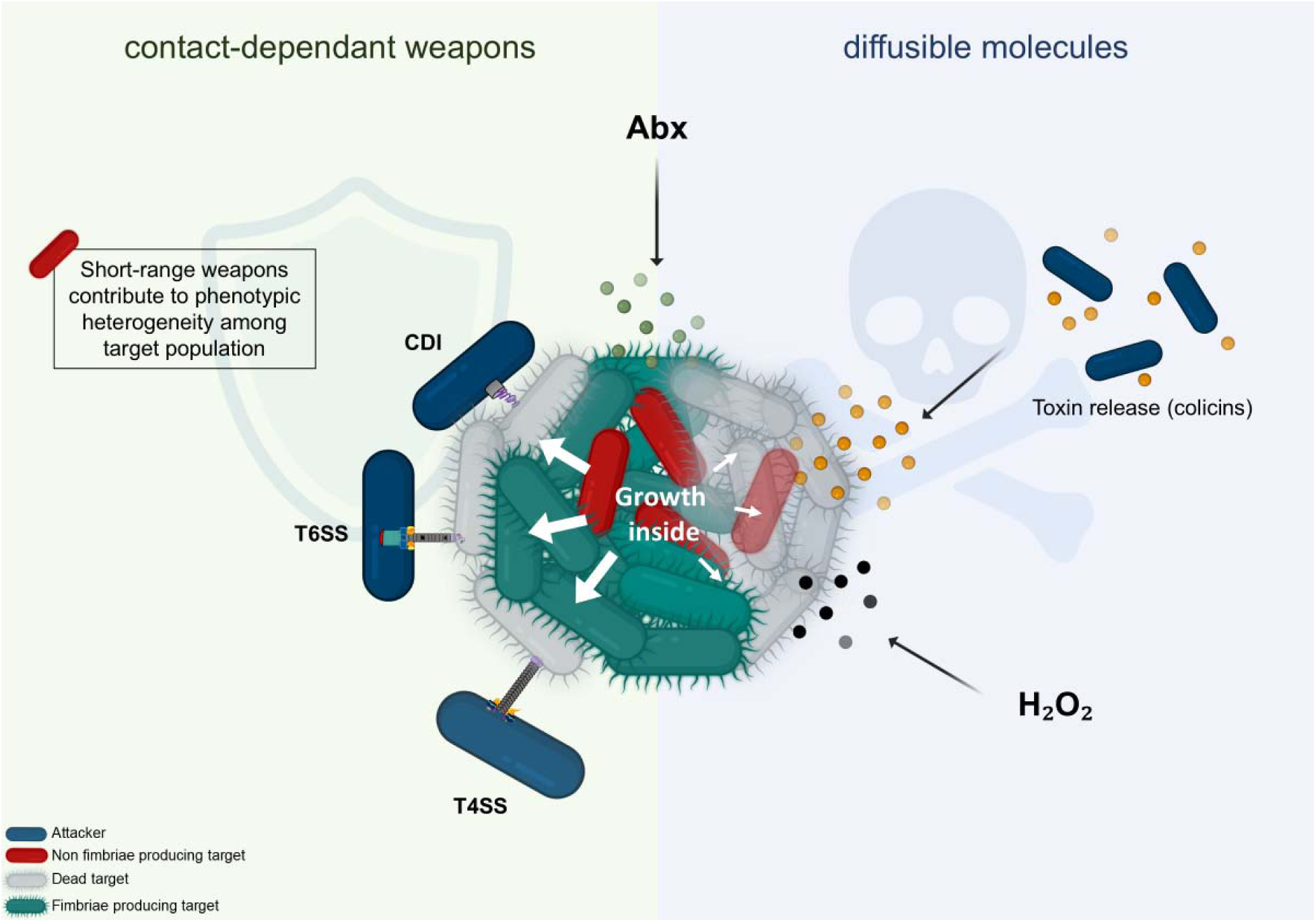
Schematic model of microcolonies resilience against diverse antibacterial weapons: the example of type 1 fimbriae mediated microcolony. **(A)** Microcolonies protect bacteria against short-range, contact-dependent weapons such as contact-dependent inhibition (CDI), type 6 secretion (T6SS) or type 4 secretion (T4SS) systems. In addition, target bacteria that do not produce type 1 fimbriae can survive inside type 1 fimbriae-mediated microcolonies supporting phenotypic heterogeneity among the target population. **(B)** Long-range, diffusible toxins such as colicins or hydrogen peroxide (H₂O₂) can fully disintegrate targeted bacterial microcolonies. Microcolonies show limited susceptibility to antibiotics (Abx).

## Materials and Methods

### Bacterial strains, media and chemicals

Bacterial strains and plasmids used in this study are listed in Supplementary Table 1. *E. coli* BW25113 WT-like, Δ*fimE*, locked-ON and locked-OFF strains, CM3267 WT, Δ*tssM, E. coli* W3110 pCH163-TreX and pCH10163 were routinely cultivated in LB-Lennox medium (5 g L^-^¹ NaCl, 5 g L^-^¹ Yeast extract, 5 g L⁻¹ tryptone, and 15 g L⁻¹ agar when needed). EAEC 17-2 WT and Δ*tssM* were grown in Sci1-inducing medium (SIM, M9 minimal medium supplemented with 1 μg mL^-^¹ of vitamin B1, 0.2 % of glycerol, 40 μg mL^-^¹ of casaminoacids and 10 % (v/v) LB medium) to induce expression of the Sci1 T6SS gene cluster*. C. rodentium* RLC55 (a ICC168 derivative in which the T6SS promoter has been swapped by a P*lac*-P*araBAD* synthetic promoter ^31^, and *L. enzymogenes* OH11 ^49^ were grown in LB at 37°C and 28°C, respectively. Expression of the *C. rodentium* T6SS gene cluster in strain RLC55 was induced with 0.5 mM of IPTG and 0.2% of L-arabinose in liquid cultures and 0.5 mM of IPTG and 2% of L-arabinose on plates ^31^. When required, media were supplemented with ampicillin (100 μg mL^-^¹), kanamycin (50 μg mL^-^¹), chloramphenicol (40 μg mL^-^¹), spectinomycin (120 μg mL^-^¹), and D-mannose 5%.

### Strains and plasmids construction

Construction of mutants were made using primers from Supplementary Table 2.

### Construction of *fimS* “locked-ON” and “locked-OFF” mutants

Locked-ON and locked-OFF mutants were constructed to lock the orientation of the *fimS* invertible promoter controlling type 1 fimbriae expression. To prevent recombinase-mediated inversion, one of the two inverted repeats (IRR) flanking *fimS* was deleted. Locked-ON and locked-OFF recombination fragments were generated using a two-step PCR strategy. In the first step, the *fimS* region was amplified in the desired ON or OFF orientation using primer pairs ON/OFF_3 with locked-ON_5 or locked-OFF_5, respectively. Locked-ON_5 and locked-OFF_5 primers were designed to anneal downstream of the IRR, thereby excluding this recombination site from the amplified *fimS* fragment. These primers also comprised homology sequences to a kanamycin (*aph*(3′)-Ia) or chloramphenicol (*cat*) resistance cassette, respectively. In the second step, overlap-extension PCR was performed to fuse the corresponding antibiotic resistance cassette, amplified from pEforCP (*aph*(3′)-Ia) or pKD3 (*cat*), to the *fimS* fragment. The resulting linear DNA fragments were introduced into *E. coli* BW25113 carrying the λ Red recombinase expressed from plasmid pSIM6, and chromosomal integration was achieved by homologous recombination^50^. Correct promoter orientation and genetic stability were verified by orientation-specific PCR (Supplementary Table 2), DNA sequencing, and electron microscopy analysis.

### Construction of the pColorSwitch reporter plasmid

The plasmid pColorSwitch was constructed using Gibson assembly. First, the pBAD30 backbone (kindly provided by Antoine Champie from the Sébastien Rodrigue lab), *rfp gene*, *fimS* promoter, and *gfp gene* fragments were amplified separately with overlapping homology regions for Gibson assembly, using oligonucleotides in Supplementary Table 2. The *rfp* fragment was designed with homology to pBAD30 and to the *fimS* promoter, such that *rfp* is positioned upstream of *fimS* and its start codon (ATG) is located immediately before the *fimS* sequence. This arrangement ensures that when the *fimS* promoter switches orientation, it drives the transcription of the *rfp* gene. The *fimS* promoter fragment was amplified with homology to the upstream *rfp* sequence and to the upstream region of *gfp*. Finally, *gfp* was amplified with homology to the upstream *fimS* sequence and to the downstream pBAD30 backbone. The four fragments were assembled using Gibson Assembly, and the resulting construct was transformed into both the WT-like and Δ*fimE E. coli* BW25113 strains. The plasmid sequence was verified by DNA sequencing.

### Interbacterial competition assay

Two different methods of interbacterial competition assay were used. The first method was based on the survivor growth kinetics (SGK) assay adapted from Taillefer *et al.* ^16^. The SGK assay is based on the lag time necessary for surviving target bacteria to reach the mid-log phase after competition (time of emergence, Te). The time of emergence directly correlates with the killing efficiency of the attacker strain on the target. The SGK method was employed for assessing the survival of *E. coli* BW25113 WT-like, Δ*fimE,* locked-ON and locked-OFF strains against CM3267, EAEC 17-2, *C. rodentium* RLC55, *L. enzymogenes* OH11, *E. coli* W3110 pCH163-TreX, and against colicins A, M and E9. Target cells, transformed with the pUC12 vector, and attackers were grown overnight in LB medium, and adjusted to an OD_600nm_ of 1. For competition with EAEC 17-2, overnight cultures of attackers and target were diluted 1/50 in SIM until OD_600nm_ reaches 0.8 before proceeding to the competition assay. Attacker and target cells were mixed in a 1:1, 4:1, 4:1, 1:1, and 1:1 ratio (v/v) for competition assays with CM3267, EAEC 17-2, *C. rodentium* RLC55, *L. enzymogenes* OH11, and *E. coli* W3110 pCH163-TreX, respectively. Then 10-µL drops of the mixture were spotted in triplicate on nitrocellulose filters deposited on pre-warmed dry LB agar plates for CM3267, *C. rodentium* RLC55, *L. enzymogenes* OH11 and *E. coli* W3110 pCH163-TreX competition assays or on SIM agar plates for competition with EAEC 17-2. Plates were incubated at 37°C for 24 h (CM3267 and *C. rodentium* RLC55) or 4 h (EAEC 17-2 and *E. coli* W3110 pCH163-TreX), and at 28°C for 8 h for *L. enzymogenes* OH11. After incubation, filters were immersed in 1 mL of LB medium supplemented with ampicillin, and cells were recovered by vortexing for 15 s. Cells were diluted 10-fold in LB supplemented with ampicillin, and 200-µL triplicates were transferred into 96-well microplates. For colicin exposure, bacterial target was adjusted to an OD_600nm_ of 1 and diluted 1/100. Then, 500 µL of diluted cultures were mixed with colicins A, M or E9 at a MOI of 1000 (1000 colicins per cell). After incubation for 30 min at 37°C, cells were harvested, washed to eliminate colicins, and resuspended into 500 µL of LB medium supplemented with ampicillin, and 200-µL duplicates were transferred into 96-well microplates. For all competition assays, growth of target cells was monitored for 15 hours at 37°C with agitation, with OD_600nm_ measurements every 5 min using a TECAN Spark 20M multimode microplate reader. The time of emergence Te was then calculated from the growth curves. The Te was normalized between the Δ*fimE*, lock-OFF, and lock-ON target strains by calculating a Te ratio, by dividing the Te of target strains by the Te of the WT-like strain. All competition assays were done at least three times.

For the second method, a plating and colony counting assay was used to evaluate competition between CM3267 wild type or mutant Δ*tssM* attackers and *E. coli* BW25113 WT-like transformed with pTrc99A, pTgH, pCaH, *E. coli* BW25113 locked-ON pRFP, locked-OFF pfuGFP or locked-ON pRFP/locked-OFF pfuGFP mix targets. Briefly, attacker and target cells were grown overnight in LB. Cultures were adjusted to an OD_600nm_ of 1, mixed in a ratio of 1:1, and 5 μL of the mixtures were spotted onto sterile nitrocellulose filter deposited on LB agar plate. After incubation for 24 h at 37°C, filters were immersed into 1 mL of LB and cells were recovered by vortexing for 15 s, serially diluted and plated onto LB selective medium. The survival of target cells was assessed using CFU counting. The survival index was calculated using the equation: 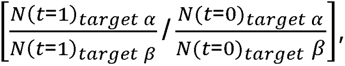, where *target α* is the target in competition with the WT attacker, *target β* the one in competition with the attacker unable to assemble it T6SS (Δ*tssM*)*, N*(*t*=1) is the numerical density in log10 at the end of the experiment and *N*(*t*=0) the one at the beginning. The experiment was repeated 3 times independently.

### Electron microscopy

A single colony of *E. coli* BW25113 wild type, locked-ON and locked-OFF strains that were grown onto agar plates at 37°C for 24h were cultured in LB liquid medium overnight. Four microliters of each culture were deposited on carbon-supported grids at room temperature. Samples were negatively stained with 2% uranyl acetate. Negatively stained bacterial cells were examined with a TECNAI G2 200 KV electron microscope (Thermo Fisher Scientific), and digital acquisitions were made with a numeric camera (Oneview, Gatan). At least 50 pictures of the three strains were taken for statistical analysis. All bacteria (complete or not) were considered for the counting of fimbriated cells.

### Growth curves

All strains were cultivated overnight at 37°C. The cultures were then adjusted to an OD_600nm_ of 1, and diluted 1/100 in 96-well microplates. Liquid plates were grown at 37°C with agitation and OD_600nm_ was measured every 20 minutes for 18 h using a TECAN Spark 20M multimode microplate reader.

### Epifluorescence microscopy

Strains to be tested were grown in liquid media overnight. Cultures were adjusted to an OD_600nm_ of 1, and 2 µl were spotted onto LB 0.8% low-melting point agar glass slides. The bacteria were observed using a Zeiss Axio Observer Z1 microscope with a Zeiss 100×/ NA Plan-Apochromat objective for figures 5B and 7A and a 60× magnification FV3000 Olympus confocal microscope for figure 1B and 8A. Images were taken either in bright light field or using GFP and RFP filters. Three independent experiments have been done. For each experiment, strains were spotted in triplicate, and three images were taken for each replicate.

### Yeast agglutination assay

*E. coli* BW25113 WT-like, locked-ON and locked-OFF strains were cultured overnight in liquid media. The cultures were adjusted to an OD_600nm_ of 6, centrifuged at 5,000 rpm for 5 min and resuspended in 5 mL of 1× PBS. D-mannose was added to bacterial culture at a concentration of 5% when required. Yeast cells were resuspended in 5 mL of 1× PBS (1%). Finally, bacteria and yeast cells were mixed with a ratio of 1:1 and 20 µL of the mixture were spotted in triplicate onto glass slides. Pictures of yeast agglutination were taken with a Google Pixel 7 camera. Experiments were performed with three biological replicates.

### Interbacterial competition assay in whole colony

Whole colony interbacterial competition assay was performed as previously described^13^. Briefly, locked-ON and locked-OFF targets, fluorescently tagged with mRFP (pRFP) and sfGFP (pfuGFP) respectively, and CM3267 WT or Δ*tssM* attackers were grown LB overnight. The cultures were then adjusted to an OD_600nm_ of 1 and mixed at a ratio of 1:1. Finally, 2 μL of the mixture were spotted on LB 0.8% noble agar in 24-well microplates and incubated at 37°C for 4 h. The co-cultures were imaged at 0 and 4 h using a 20× magnification FV3000 Olympus confocal microscope. The assay was independently performed twice, with three technical replicates.

Due to technical constraints in this assay (and in the interbacterial competition assay shown in Figure 6C), locked-ON cells were tagged with mRFP and locked-OFF cells with sfGFP. However, for consistency with the rest of the figures in the manuscript, we intentionally inverted the color representation during image processing. Locked-ON cells are therefore shown in green, and locked-OFF cells in red.

### Antibiotics and hydrogen peroxide survival assay

Target bacteria resistance against different antibiotics was evaluated as follows. Cells were grown overnight in LB medium and adjusted to an OD_600nm_ of 1. Fifty microliters of each culture were mixed with 50 μL of LB containing either no antibiotic or twice the final concentration of one of the following antibiotics: ciprofloxacin (0.25 μg mL^-^¹), tobramycin (16 μg mL^-^¹), gentamicin (8 μg mL^-^¹) or ceftriaxone (16 μg mL^-^¹). Mixtures were transferred in triplicate into 96-well microplates. For the hydrogen peroxide (H₂O₂) survival assay, LB medium was supplemented with either 0.03125 M or 0.0625 M H₂O₂. Plates were then incubated for 90 min at 37°C for antibiotics assays and 20 min at room temperature for H₂O₂ treatment. After incubation, 10 μL of each mixture were serially diluted, and 3 μL were spotted onto LB agar plates for CFU enumeration. The survival index was calculated following this equation: 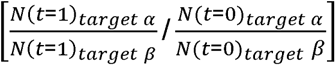 in which *target α* corresponds to cells exposed to antibiotics or H₂O₂, *target β* corresponds to untreated controls, *N*(*t*=1) is the numerical density (log₁₀ CFU) at the end of the experiment, and *N*(*t*=0) the density at the beginning. The experiment was repeated 3 times independently. Survival of target strains was determined by colony-forming unit (CFU) counts. For samples in which no colonies were observed, the assay’s limit of detection (dl) was assigned as the minimum detectable value.

### MIC determination

Minimum inhibitory concentrations inhibiting 90% of bacterial growth (MIC) were determined. *E. coli* BW25113 WT-like, Δ*fimE*, locked-ON, and locked-OFF strains were grown overnight in LB medium at 37 °C. Cultures were adjusted to an OD_600nm_ of 1 and 1:100 of cultures was then used to inoculate 96-well microplates containing two-fold serial dilutions (64 to 0.015625 µg mL^-^¹) of ciprofloxacin, delafloxacin, ceftriaxone, gentamicin, or tobramycin. Each condition was tested in triplicate. Plates were incubated at 37 °C for 24 h, after which bacterial growth was measured by monitoring OD_600nm_ using a TECAN microplate reader. MIC values were defined as the lowest antibiotic concentration resulting in ≥ 90% inhibition of growth compared to the no-antibiotic control for each strain.

### *fimS* switching colony assay

E. coli BW25113 WT-like, WT-like pColorSwitch, Δ*fimE* pColorSwitch, locked-ON pRFP, and locked-OFF pfuGFP strains were grown overnight in LB medium at 37 °C with shaking, supplemented with ampicillin (100 µg mL^-^¹), kanamycin (50 µg mL^-^¹), or chloramphenicol (50 µg mL^-^¹) when required. Cultures were normalized to an OD_600nm_ of 1, and serial dilutions were prepared in 96-well plates. For each strain, 50 µL of the 10⁻□ dilution was spread onto LB agar plates containing the appropriate antibiotics. After incubation, isolated colonies were imaged using a Vilber imaging module equipped with GFP and RFP emission filters. GFP⁺, RFP⁺, and dual GFP⁺/RFP⁺ colonies were quantified and analyzed using CellProfiler. Experiments were performed with three biological replicates, each including three technical replicates.

### Statistical analysis of *fimS* switching

SfGFP and mRFP fluorescence images were first combined using CellProfiler (v4.2.8) ^51^ to create a single composite image containing all colonies. A mask corresponding to the interior of the Petri dish was generated restrict subsequent analyses to the valid imaging area. Combined images were then used as input in Cellpose (v4) ^52^ to produce segmentation masks. These segmentation masks were subsequently imported into CellProfiler, where fluorescence intensity measurements were extracted from the GFP and RFP channels, enabling classification of colonies as GFP-positive, RFP-positive, or double-positive.

### Statistical analysis

All experiments were carried out at least three times with at least 3 technical replicates for each experiment except for the electron microscopy assay and the whole colony interbacterial competition assay. Statistical analyses were made using the GraphPadPrism 9 software. A threshold of *p* < 0.01 was used to define statistical significance.

## Acknowledgements

We thank Daniel Garneau, M.Sc., for his excellent technical assistance with confocal and epifluorescence microscopy, as well as with data analysis using Cellpose and CellProfiler. This work was supported by the Natural Science and Engineering Research Council of Canada (RGPIN-2019-06044). JPC holds a Chercheur-Boursier junior 2 award from the Fonds de recherche du Québec-Santé. Work on T6SS in the Cascales lab is supported by the Centre National de la Recherche Scientifique, the Aix-Marseille Université, and by grants from the Agence Nationale de la Recherche (ANR-20-CE11-0017 and ANR-24-CE44-7545) and the Fondation Bettencourt-Schueller. MMD is supported by a doctoral fellowship from the Fonds de recherche du Québec-Nature et technologies.

## Author contributions

MMD, EC and JPC conceived the study and designed the experiments. MMD, JFG, AK and GQ contributed to the investigation. GQ provided bacterial strains. MMD, JPC and EC analyzed the data and wrote the manuscript. All authors read, revised the manuscript and approved the final version.

