## Supplementary material for "Aggregation-mediated microcolony shields bacteria from contact-dependent competition and maintains phenotypic heterogeneity": suppl. Tables

- **Tables S1-S2**
- **References**

**Supplementary Tables**

**Table S1. Bacterial strains and plasmids used in this study.**

| **Bacterial strains** | **Reference** |
| --- | --- |
| *Cronobacter malonaticus* 3267 | ^1^ |
| *Cronobacter malonaticus* 3267 Δ*tssM::aadA(SmR)* | ^1^ |
| *E. coli* BW25113 Δ*yejO::nptII(KanR)* | ^2^ |
| *E. coli* BW25113 Δ*fimE::nptII(KanR)* | ^2^ |
| *E. coli* BW25113 locked-ON | This study |
| *E. coli* BW25113 locked-OFF | This study |
| EAEC 17-2 | Arlette Darfeuille-Michaud |
| EAEC 17-2 Δ*tssM* | ^3^ |
| *Lysobacter enzymogenes* OH11 | ^4^ |
| *Lysobacter enzymogenes* OH11 Δ*virD4* | ^4^ |
| *Citrobacter rodentium* RLC55 | ^5^ |
| *Citrobacter rodentium* RLC55 Δ*tssM* | ^5^ |
| *E. coli* W3110 | ^6^ |
| **Plasmids** | **Reference** |
| pCH10163 | ^7^ |
| pCH10163-TreX | ^7^ |
| pUC12 | ^8^ |
| pTrc99A | Pharmacia Biotech |
| pTghH | ^9^ |
| pCah | ^10^ |
| pBAD30 | ^11^ |
| pColorSwitch | This study |
| pKD3 | ^12^ |
| pSIM19 | ^13^ |
| pSIM6 | ^14^ |
| pfuGFP | New York Botanics, LLC |
| pRFP | ^10^ |

**Table S2. Primers used in this study (5’ to 3’).**

Primers for locked-ON/ locked-OFF construction:

**Amplification of *fimS* deleted of the IRR region**

ON/OFF_3

GCTGCTTTCCTTTCAAAAAACTATTT

Locked-ON_5

GAATAAATTGCAGTTTCATTTGATGCTCGATGAGTTTTTCTAAGAATTCTTTTGACTCATAGAGGAAAGCATCGCGGAC

Locked-OFF_5

TTTCATTATGGTGAAAGTTGGAACCTCTTACGTGCCGATCAGGCCAAACTGTCCATATCATAAATAAGTTACGTATT

**Amplification of *aph(3')-Ia*-fimS-locked-ON**

ON_5

AACTTATTGATAATAAAGTTAAAAAAACAAATAAATACAAAAGCCACGTTGTGTCTCAAAAT

**Amplification of *cat*-fimS-locked-OFF**

OFF_5

AACTTATTGATAATAAAGTTAAAAAAACAAATAAATACAAGGAACTTCATTTAAATGGCGCGCCT

**PCR Verification**

Int_ON_3_OFF_5

TGTCCGCGATGCTTTCCTCTA

Ext_ON_5

GTGAGTAAACGTCGTTATCTTACCG

Ext_OFF_3

GAACCTGGGTAGGTTATTGATACTGAA

Primers for pColorSwitch construction:

**Amplification of pBAD30**

pBAD30_F

GGATCCTCTAGAGTCGACCTGCAG

pBAD30_R

GGTACCGAGCTCGAATTCGCT

**Amplification of RFP**

Overlap-pBAD30_RFP _5

ATACCCGTTTTTTTGGGCTAGCGAATTCGAGCTCGGTACCTCTACTTGTACAGCTCGTCCAT

RFP_overlap-*fimS*_3

ATTTATGATATGGACAGTTTGGCCCCAATTGTCTTGTATTGAATTCGAGCTCGGTACCCG

**Amplification of fuGFP**

fuGFP_overlap-pBAD30 _3

AGCCAAGCTTGCATGCCTGCAGGTCGACTCTAGAGGATCCcttattagctcttatacagctcgtccat

Overlap-*fimS*_fuGFP_5

CCATGTCGATTTAGAAATAGTTTTTTGAAAGGAAAGCAGCatggtgagcagtggcgaag

**PCR Verification**

Int_RFP_3

GAGGGCTTCAAGTGGGAGC

Int_fuGFP_3

CGAGTTTTGTACGTGCCATCTTCC

Ext_pBAD30_R

CCGCCAGGCAAATTCTGTTT

Ext_pBAD30_F

CGGCGTCACACTTTGCTATG
